# Chronic Jetlag Accelerates Pancreatic Neoplasia in Conditional *Kras*-Mutant Mice

**DOI:** 10.1101/2022.07.18.500370

**Authors:** Patrick B Schwartz, Morgan T Walcheck, Manabu Nukaya, Derek M Pavelec, Kristina A Matkowskyj, Sean M Ronnekleiv-Kelly

## Abstract

Misalignment of the circadian clock compared to environmental cues causes circadian desynchrony, which is pervasive in humans. Clock misalignment can lead to various pathologies including obesity and diabetes, both of which are associated with pancreatic ductal adenocarcinoma - a devastating cancer with an 80% five-year mortality rate. Although circadian desynchrony is associated with an increased risk of several solid-organ cancers, the correlation between clock misalignment and pancreas cancer is unclear. Using a chronic jetlag model, we investigated the impact of clock misalignment on pancreas cancer initiation in mice harboring a pancreas-specific activated *Kras* mutation. We found that chronic jetlag accelerated the development of pancreatic cancer precursor lesions, with a concomitant increase in precursor lesion grade. Cell-autonomous knock-out of the clock in pancreatic epithelial cells of *Kras*-mutant mice demonstrated no acceleration of precursor lesion formation, indicating non-cell-autonomous clock dysfunction was responsible for the expedited tumor development. Therefore, we applied single-cell RNA sequencing over time and identified fibroblasts as the cell population manifesting the greatest clock-dependent changes, with enrichment of specific cancer-associated fibroblast pathways due to circadian misalignment. Collectively, these results suggest fibroblasts as the putative target of chronic jetlag-induced accelerated pancreas cancer initiation.

## Introduction

Circadian clock proteins coordinate daily biological rhythms with an oscillation of 24 hours (McIntosh et al. 2010). Control from the central clock in the hypothalamus assimilates environmental cues (e.g. day/night cycles) to modulate peripheral organ rhythmic expression of core circadian genes and behavioral patterns (e.g. sleep/wake cycle) (McIntosh et al. 2010). In addition to central control, nearly all cells have an autonomous circadian clock based on transcriptional and translational feedback mechanisms of core clock genes (CCGs) (Lee et al. 2001; Preitner et al. 2002; Sato et al. 2004; Takahashi 2017). Collectively, the CCGs of the central and peripheral clocks orchestrate the transcription of circadian output genes, also known as clock-controlled genes (Takahashi 2017; Cox and Takahashi 2019). The central and peripheral clocks in mammals (and other organisms) have evolved to anticipate environmental changes that occur over the 24-hr day/night cycle, and in doing so, appropriately sequence complex internal processes to align with external cues (Takahashi 2017). For instance, in humans, energy intake, physical activity, and cognitive activities are anticipated to occur during the day while rest and sleep are expected to occur at night (Pittendrigh 1993; Wright et al. 2013). Consequently, this network of circadian control coordinates the optimal timing of critical cell and organ processes such as metabolism, cell division, apoptosis, and immune function (Matsuo et al. 2003; Bass and Takahashi 2010; Lee and Sancar 2011; Scheiermann et al. 2013).

Notably, the internal clock can be adjusted, or re-entrained, based on changes to environmental zeitgebers or ‘time-givers’, such as through alteration of the day/night cycle (Wright et al. 2013). However, persistent misalignment between environmental cues compared to the internal clocks can result in circadian desynchrony with associated aberrant gene signaling (Kettner et al. 2016; West et al. 2017). Unfortunately, this type of clock dysfunction (misalignment or circadian desynchrony) is pervasive in humans and this condition affects a substantial proportion of the human population, particularly shift workers, whose clock timing can be altered dramatically compared to environmental zeitgebers by work schedule requirements (Bass and Lazar 2016). Further, this environmental or behavior-induced desynchrony can have profound implications for human health including the promotion of various pathologies such as the metabolic syndromes of obesity and diabetes (Arble et al. 2009; Roenneberg et al. 2012; Perelis et al. 2016). There is also a strong correlation between circadian clock dysfunction and the risk of developing several different solid organ cancers, including lung cancer, liver cancer, and colon cancer (Schernhammer et al. 2003; Papagiannakopoulos et al. 2016; Kettner et al. 2016). Meanwhile, a history of shiftwork has been shown to increase the risk of pancreatic ductal adenocarcinoma (PDAC) in men over 2-fold (Parent et al. 2012); concordantly, obesity and diabetes are both risk factors for the development of PDAC (Schernhammer et al. 2003; Marcheva et al. 2010; Kettner et al. 2016). Despite this converging evidence and the substantive data linking the circadian clock to pancreas-specific disease processes (Marcheva et al. 2010; Perelis et al. 2015), the correlation between circadian desynchrony and the risk of developing PDAC remains unclear.

We sought to establish the connection between clock misalignment – a process that is highly prevalent in humans – and PDAC because this would have important implications for understanding cancer pathogenesis and risk mitigation in this deadly disease. Risk factor reduction leading to cancer prevention is particularly relevant in this highly lethal malignancy because few people survive after cancer diagnosis: of the 60,430 people estimated to be diagnosed in 2021, approximately 48,220 (∼80%) will die of their disease (Siegel et al. 2021). As a foundation for uncovering the link between circadian clock dysfunction and the development of PDAC, we investigated whether circadian misalignment impacted tumor initiation within the pancreas. We used *LSL*-*Kras*^*G12D/+*^; *Pdx1-Cre* mice (KC) that harbor a *Kras* mutation in pancreatic acinar and ductal cells (Hingorani et al. 2003). KC mice develop the full spectrum of neoplastic changes including acinar-to-ductal metaplasia (earliest lesion), pancreatic intra-epithelial neoplasia 1 (PanIN-1), higher grade PanIN lesions (PanIN-2, -3), and ultimately PDAC (Hingorani et al. 2003). We implemented a well-characterized model of chronic jetlag (CJ) that is known to cause environmental-induced misalignment and phase-shift of the clock (Filipski et al. 2004; Papagiannakopoulos et al. 2016; Schwartz et al. 2021) and is meant to mimic the environmental and behavioral misalignment experienced by humans (Vetter 2020). The abrupt change in timing of the light/dark cycle with CJ causes a period of desynchrony, with differences in the rate of re-entrainment by internal clocks or behavior (Yamazaki et al. 2000; West and Bechtold 2015). Previously, we have found that behavioral re-entrainment occurs much more rapidly than re-entrainment of the pancreatic clock (Schwartz et al. 2021). Moreover, this type of protocol (CJ) has been shown to promote the growth of several other cancers, including osteosarcoma, and hepatocellular carcinoma in mice (Filipski et al. 2004; Filipski et al. 2009; Kettner et al. 2016).

In the current study, we hypothesized that circadian clock desynchrony through CJ would accelerate the initiation of PanIN-1 and progression to higher-grade PanINs or PDAC. We found that the pancreata of KC mice subjected to CJ developed increased fibroinflammatory infiltrate and neoplastic lesion formation (i.e. PanINs) compared to KC mice under standard conditions. To understand whether cell-autonomous clock dysfunction was responsible for the accelerated PanIN development, we generated KC mice with an abolished clock (*Bmal1* knock-out) in the pancreatic ductal and acinar cells (cellular origin of PDAC). These mice did not replicate the phenotype observed in the CJ group of KC mice, indicating a non-cell-autonomous role of clock dysfunction that is facilitating the PanIN progression. To investigate further, we used single-cell RNA sequencing (scRNA-seq) and identified cancer-associated fibroblasts (CAFs) as the putative target of the CJ-induced phenotype. This work serves as a foundation for understanding the link between circadian misalignment through CJ and the acceleration of PDAC development.

## Materials and Methods

### Mouse Care and Husbandry

All animal studies were conducted according to an approved protocol (M005959) by the University of Wisconsin School of Medicine and Public Health (UW SMPH) Institutional Animal Care and Use Committee (IACUC). Male and female mice were housed in an Assessment and Accreditation of Laboratory Animal Care (AALAC) accredited selective pathogen-free facility (UW Medical Sciences Center) on corncob bedding with chow diet (Mouse diet 9F 5020; PMI Nutrition International) and water *ad libitum*.

Conditional *LSL-Kras*^G12D/+^ (K) mice (B6.129S4-Kras tm4Tyj/J (#008179)) were bred with mice expressing Cre-Recombinase driven by the *Pdx1* promoter (C) (B6.FVB-Tg(Pdx1-cre)6Tuv/J) (#014647)) to generate *LSL-Kras*^G12D/+^, *Pdx1*-*Cre* (KC) mice (Hingorani et al. 2003). The KC mouse has been well characterized as a pancreas pre-cancer model (Hingorani et al. 2003). To evaluate the effect of loss of clock function on KC mouse pathology the conditional *Bmal1*^*fx/fx*^ mouse, kindly gifted by Professor Christopher Bradfield (University of Wisconsin-Madison, Madison, WI), was crossed with the KC mouse to generate the *Bmal1*^*fx/fx*^, *LSL-Kras*^G12D/+^; *Pdx1*-*Cre* (BKC) mouse model (Liu et al. 2014; Johnson et al. 2014). To identify cell types targeted by *Pdx1*-*Cre* recombinase in the KC mouse model, the KC mouse was additionally crossed with the *LSL-tdTomato* reporter (Ai14) mouse (B6.Cg-Gt(ROSA)26Sortm14(CAG-tdTomato)Hze/J (#007914)) to produce the KCT mouse (*LSL-R26*^*CAG-tdTomato*^; *LSL-Kras*^G12D/+^; *Pdx1*-*Cre*). The presence of each allele, including activated *Kras*-mutation, was confirmed with genotyping PCR according to Jackson Laboratory’s (Bar Harbor, ME) protocols as previously described (Liu et al. 2014; Walcheck et al. 2021). The K, C, and T mice were purchased from Jackson Laboratory.

### Chronic Jetlag Protocol

Starting at 4-6 weeks of age, KC mice were subjected to either a normal circadian (standard lighting) 12-hour/12-hour light: dark (LD) cycle condition, or a chronic jetlag protocol (CJ) known to induce circadian desynchrony and mimic altered environmental exposure experienced by humans (Filipski et al. 2004; Papagiannakopoulos et al. 2016; Vetter 2020; Schwartz et al. 2021). CJ consisted of a 12-hour/12-hour LD cycle, phase-shifted forward 4 hours every 2-3 days. Mice were kept under their respective conditions until the study endpoint at 5 and 9 months of age.

### Histologic Analysis

Mouse pancreatic tissue was fixed in 10% neutral buffered formalin for 48 hours and then stored in 70% ethanol until further processing. Tissue was processed, paraffin embedded, sectioned onto slides, and stained with hematoxylin and eosin (H&E) by the University of Wisconsin Carbone Cancer Center (UWCCC) Experimental Animal Pathology Laboratory (EAPL) Core. Each pancreas was sectioned at 200 μm steps for at least 6 slides. All slides processed were assessed for the presence of pancreas cancer precursor lesions (pancreatic intraepithelial neoplasia or PanIN) along with grading on a scale of 1-3 (1 = lowest grade, 2 = intermediate grade, and 3 = high grade) and/or pancreatic ductal adenocarcinoma (PDAC). This was performed by a blinded, board-certified, gastrointestinal pathologist (KAM) at 5 months (n = 36 normal circadian [24 male and 12 female] vs n = 26 CJ [14 male and 12 female]) and 9 months age (n = 33 normal circadian [19 male and 14 female] vs n = 39 CJ [23 male and 16 female]). Additional BKC mice (n = 24 [14 male and 10 female]) were assessed at 5 months. Categorical comparisons between groups were made with the Chi-squared or Fisher’s Exact test as appropriate. KC mice develop fibroinflammatory infiltrate associated with adjacent PanINs and are collectively part of the pancreatic changes with neoplasm formation (Hingorani et al. 2003). Quantification of the total percent pancreas involvement by the fibroinflammatory infiltrate and PanINs (referred to collectively as FI/PanIN) was made by 3 separate reviewers, and differences were tested using Student’s t-test between groups. For evaluation, two slides were selected from each pancreas comprising the largest area of available tissue. Values are presented as the mean with associated standard deviation (SD). Inter-rater reliability statistics were computed and the intraclass correlation coefficient is reported with associated 95% confidence intervals.

### Immunohistochemistry

KCT mouse pancreas samples were formalin-fixed, paraffin embedded, and sectioned by the EAPL Core facility. Immunohistochemistry (IHC) for red fluorescence protein (tdTomato) was also performed by EAPL. Sections were deparaffinized in xylenes and hydrated through graded alcohols to distilled water. Antigens were retrieved using citrate buffer pH 6.0 (10 mM Citric Acid and 0.05% tween 20).

Endogenous peroxidase was blocked with 0.3% H_2_O_2_ and blocking of non-specific binding was performed using 10% goat serum. Sections were incubated with rabbit anti-RFP antibody (600-401-379, Rockland Inc, Pottstown, PA) (1:1600) followed by ImmPRESS goat anti-rabbit IgG (MP-7451, Vector Laboratories, Burlingame, CA). Detection was performed using the DAB substrate kit (8059S, Cell Signaling Technology, Danvers, MA). Samples were counterstained using Mayer’s hematoxylin (MHS32, Millipore-Sigma, St. Louis, MO).

### scRNAseq Sample Preparation, Library Construction, and Sequencing

Five- and 9-month-old normal circadian and CJ KC mice were sacrificed with cervical dislocation and the pancreas was rapidly dissected (< 60 seconds) and placed in cold PBS (Thermo Fisher Scientific, Waltham, MA). One male and female pancreas was pooled from each condition and time point after demonstration of no discernable sex-specific differences (Takele Assefa et al. 2020). Pancreas samples were minced until the tissue was < 1 mm in diameter, and then centrifuged at 900 RPM for 5 minutes to remove fat contamination. Samples were then incubated with 5 mL of Collagenase P (Sigma Aldrich, St. Louis, MO) in HBSS (Thermo Fisher Scientific) at 37° C for 15 minutes while being shaken at 100 RPM. Additionally, samples were manually agitated every 2-3 minutes. The digestion reaction was quenched with 10 mL R10 media. Samples were then centrifuged at 900 RPM and washed twice with 10 mL of R10. Samples were then strained in a gradient fashion using 500 μm, 100 μm, and 40 μm strainers. Samples were then live/dead sorted into single-cell populations and were transferred to the University of Wisconsin Gene Expression Center (GEC) for Library Preparation.

In brief, libraries were constructed according to the Chromium NextGEM Single Cell 3’ Reagent Kit v3.1 User Guide, Rev.D (10x Genomics, Pleasanton, CA). Single-cell suspension cell concentration and viability were quantified on the Countess II (Thermo Fisher Scientific) using 0.4% Trypan Blue (Invitrogen, Carlsbad, CA). Single-cell droplet encapsulation was performed using the Single Cell G Chip and the 10x Genomics Chromium Controller. Following the completion of the Chromium run, the Gel Beads in Emulsion (GEMs) were transferred to emulsion-safe strip tubes for GEM-RT using an Eppendorf MasterCycler Pro thermocycler (Eppendorf, Hamburg, Germany). Following reverse transcription (RT), GEMs were broken, and the pooled single-cell cDNA was collected in Recovery Agent, purified using DynaBeads MyOne Silane beads (Thermo Fisher Scientific), and amplified with the appropriate number of PCR cycles based on the number of cells targeted for recovery. Post-cDNA amplified product was purified using SPRIselect (Beckman Coulter, Brea, CA) and quantified on a Bioanalyzer 2100 (Agilent, Santa Clara, CA) using the High Sensitivity DNA kit. Adapters were then added to the libraries after fragmentation, end repair, A-tailing, and double-sided size selection using SPRIselect. Following adapter ligation, libraries were cleaned up using SPRIselect and sample-specific indexes (Chromium i7 Multiplex Kit, 10x Genomics) were added by sample index PCR using the appropriate number of PCR cycles based on the cDNA yield calculated by the Bioanalyzer quantification. After sample index PCR, samples were double-size selected using SPRIselect and quantified by the Qubit High Sensitivity DNA Kit (Thermo Fisher Scientific). Libraries were sequenced on a NovaSeq6000 (Illumina, San Diego, CA).

### scRNAseq Data Analysis

Single-cell RNAseq data were analyzed by the UW Bioinformatics Resource Center. Data were demultiplexed using the Cell Ranger Single Cell Software Suite, mkfastq command wrapped around Illumina’s bcl2fastq. The MiSeq balancing run was quality-controlled using calculations based on UMI-tools (Smith et al. 2017). Sample libraries were balanced for the number of estimated reads per cell and run on an Illumina NovaSeq system. Cell Ranger software was then used to perform demultiplexing, alignment, filtering, barcode counting, UMI counting, and gene expression estimation for each sample according to the 10x Genomics documentation (https://support.10xgenomics.com/single-cell-gene-expression/software/pipelines/latest/what-is-cell-ranger). The gene expression estimates from each sample were then aggregated using Cellranger (cellranger aggr) to compare experimental groups with normalized sequencing-depth and expression data.

Gene expression data was then processed using the Seurat version 4 pipeline for data integration (Hao et al. 2021). Data was loaded into Seurat. Low quality (<200 features or >20% mitochondrial gene content) and doublets were removed. Cells expressing more than 1% hemoglobin gene content were filtered out. The normal circadian and CJ KC data were merged and integrated using the SCTransform v2 wrapper with 3000 unique features (Choudhary and Satija 2022). SCTransform allows for normalization and better batch correction based on a regularized negative binomial regression. The regression additionally allowed for correction of the cell cycle effect and mitochondrial gene content (Nestorowa et al. 2016). A principal component analysis (PCA) was performed with the number of dimensions (principal components) selected based on an elbow plot. Data were visualized using the Uniform Manifold Approximation and Projection for Dimension Reduction (UMAP) (McInnes et al. 2020). Cell identities were called based on the expression of known markers for each cell type and scored using the ScType package to cross-validate identities following the default settings for SCT-transformed data in the provided vignette (Ianevski et al. 2022). Differential expression between conditions and time points was performed between clusters or conditions in Seurat using the MAST test based on a log fold change threshold of 0.25 and a false discovery rate (FDR) corrected *p* value of *p* < 0.05. The MAST test within Seurat is based on the MAST package, which is specifically designed to handle differential expression in scRNA-seq data and utilizes a hurdle model (Finak et al. 2015). Gene Set Enrichment Analysis (GSEA) was conducted with the fgsea package leveraged against the oncogenic signature gene set (c6.all.v7.5.1.symbols) and biologic process subset of gene ontology (GO BP) (c5.go.v7.5.1.symbols) (Sergushichev 2016). Pseudotime trajectory analysis was performed with the Monocle 3 package (Trapnell et al. 2014). Genes correlated with pseudotime progression were computed using the Moran’s I test and those genes with an FDR *p* < 0.05 were considered significant. All scRNAseq data is publicly available through gene expression omnibus (GEO) (Accession number: GSE209629). All analyses in this manuscript were conducted in R version 4.2.0 (Vienna, Austria) or GraphPad Prism version 8.3.1 (Dotmatics, Boston, MA).

## Results

### Chronic Jetlag Promotes Pancreatic Tumor Initiation

To test the hypothesis that circadian misalignment by chronic jetlag (CJ) accelerates tumor initiation (i.e. PanIN or pre-cancerous lesion formation) and development of PDAC, we elected to utilize the *LSL-Kras*^G12D/+^, *Pdx1*-*Cre* (KC) mouse model – a well-characterized PDAC development model – and assessed the burden of neoplastic lesions (i.e. PanIN) and PDAC (Hingorani et al. 2003). We have previously shown that the CJ protocol results in circadian misalignment with phase-shift of core clock gene (CCG) expression and alteration of diurnally expressed genes in the pancreas compared to mice under normal circadian (standard lighting) conditions (Schwartz et al. 2021). Furthermore, we have demonstrated that after normalizing the time of lights on/lights off (i.e. light/dark cycle) in CJ mice, complete re-entrainment of the pancreatic clock does not occur out to day 10 post-normalization; this is in contrast to behavioral re-entrainment which occurred by 3 days (Schwartz et al. 2021). Therefore, the CJ protocol is a robust protocol to evaluate the impact of circadian misalignment on the tumor initiation and progression of pancreatic pre-cancerous lesions.

We first evaluated KC mice at age 5 months, because existing literature demonstrates that 5-month KC mice develop neoplastic changes consisting of early PanINs (i.e. PanIN-1) but most of the pancreas remains normal (Hingorani et al. 2003). Therefore, evaluating the extent of pancreas involvement by PanIN at this time point (not incidence) should allow for the detection of any acceleration in tumor initiation due to circadian misalignment (CJ). All sections were scored for their degree of pancreatic involvement by neoplastic lesions (e.g. PanINs) and the associated fibroinflammatory infiltrate, hereafter referred to as FI/PanIN. This was calculated by determining the percent of the pancreas that was involved with FI/PanIN by 3 independent reviewers, and there was an excellent intraclass correlation coefficient of 0.896 (95% CI 0.868-0.919; *p* < 0.01). We found that KC mice that were exposed to CJ (n = 26) displayed an approximate 2-fold mean increase (± standard deviation [SD]) in FI/PanIN involvement of the pancreas at 5 months compared to KC mice under standard lighting conditions (n = 36) (22.47% ± 19.08 vs 12.73% ± 12.08; *p* = 0.027, **Figure 1**). There were no sex-specific differences at 5 months for normal circadian (males 13.14% ± 12.68 vs. females 11.93% ± 11.27; *p* = 0.77) or CJ KC mice (males 17.27% ± 12.94 vs. females 28.53% ± 23.55; *p* = 0.16).

**Figure 1:**
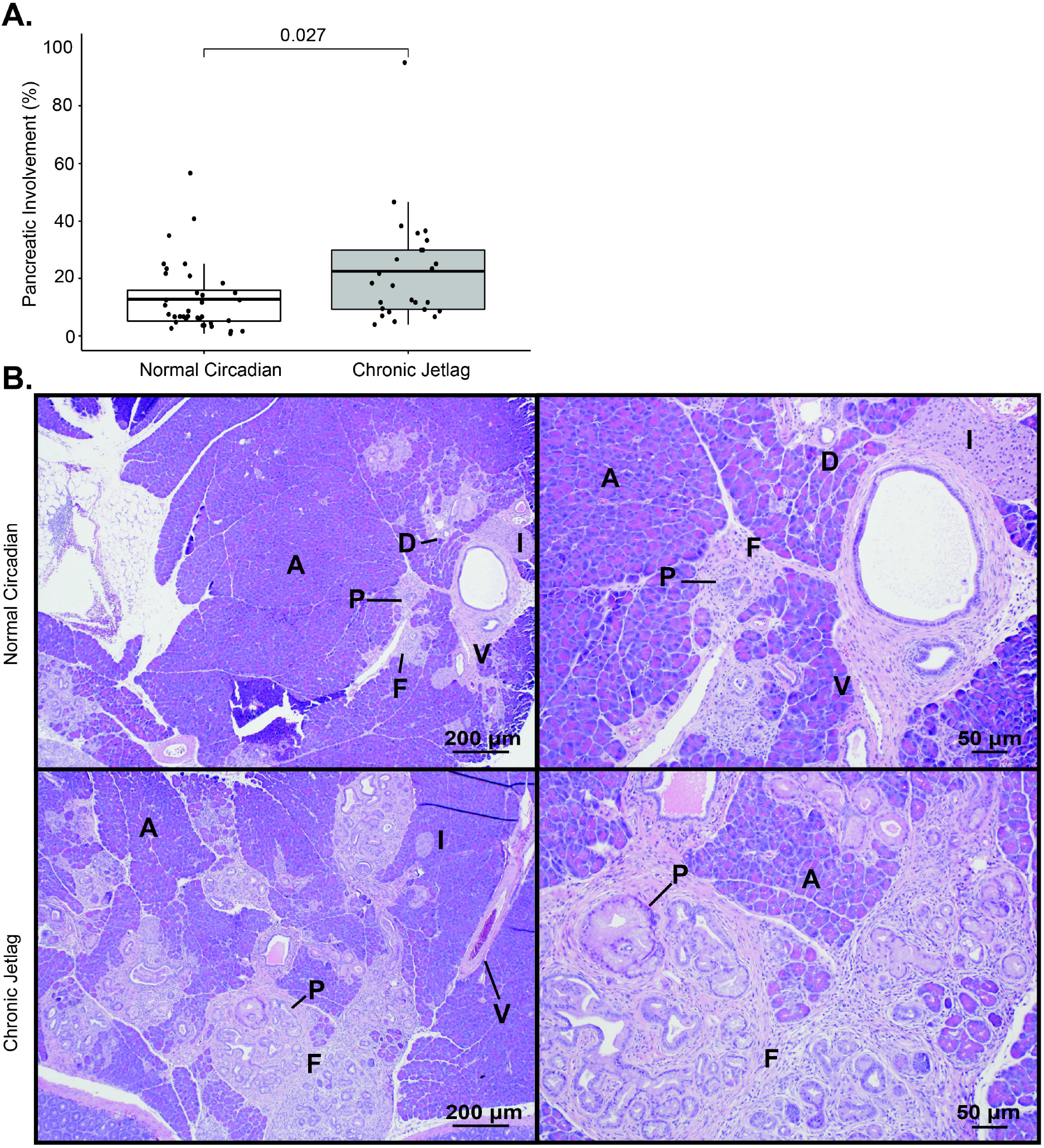
Pancreatic fibroinflammation/PanIN development is increased by chronic jetlag at 5 months. **A**. Boxplot comparing the mean (with associated 25^th^ and 75^th^ quantiles) percent pancreatic involvement of FI/PanIN at 5 months for normal circadian KC (n= 36) and CJ KC mice (n = 26). **B**. Representative H&E images of normal circadian (upper) and chronic jetlag (lower) mice at 40x (left) and 100x (right). Scale bars are shown for reference. [Annotation: PanIN (P), acinar (A) cells, islet (I) cells, ductal (D) cells, vascular (V) cells (i.e. endothelial cells), and fibroinflammatory (F) infiltrate]

Due to the expected progression of FI/PanIN at 9 months, the extent becomes less relevant (Hingorani et al. 2003). Instead, by 9 months, the incidence of higher grade PanINs (i.e. PanIN-2 and PanIN-3) and development of PDAC better reflects the progression of pre-cancerous lesions into cancer. This is because i) not all PanIN-1s progress to higher-grade lesions, ii) PanIN-2,-3, and PDAC represent tumor progression, and iii) KC mice are expected to develop a significant incidence of PanIN-2, -3, and PDAC with increasing age (Hingorani et al. 2003). In contrast, higher-grade lesions are rarely observed by approximately 5 months of age (Hingorani et al. 2003). Concordantly, we found that KC mice under normal circadian conditions exhibited almost exclusively PanIN-1, with only 5.6% (n = 2/36) demonstrating a PanIN-2 lesion at age 5 months; no 5-month KC mice under normal conditions developed PanIN-3 or PDAC (**Figure 2A/C**). Interestingly, although not significant, 7.7% (n = 2/26) of KC mice subjected to CJ developed PanIN-2, and one mouse developed dedifferentiated PDAC which had replaced the entirety of the pancreatic parenchyma (**Figure 2A/C**).

**Figure 2:**
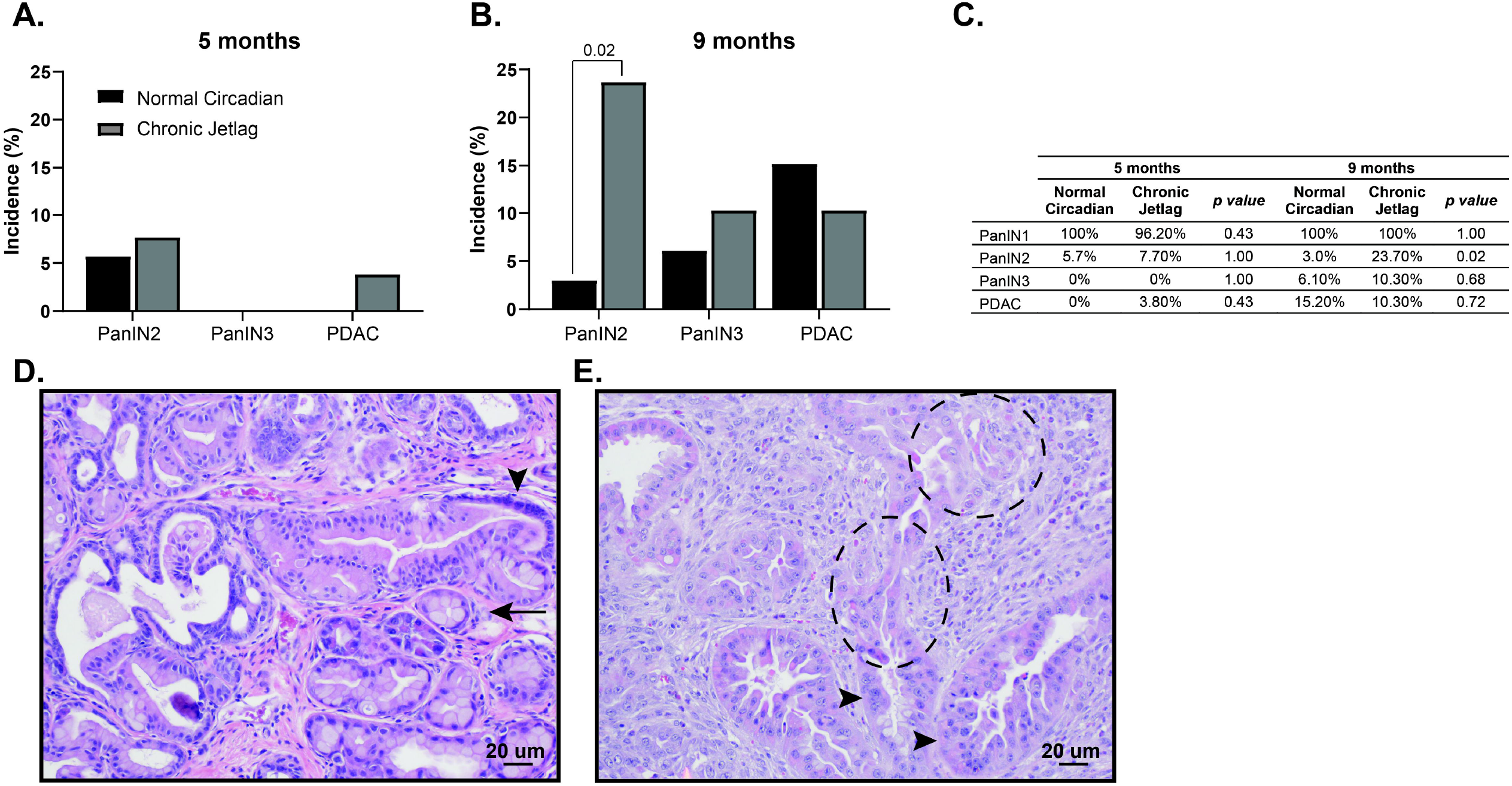
Chronic jetlag accelerates the grade of PanIN lesions. Bar charts indicating the incidence of PanIN-1-3 and PDAC in normal circadian (black) and CJ (grey) conditions at **A**. 5 months (normal circadian n = 36 vs. CJ n = 26) and **B**. 9 months (normal circadian n = 33 vs. CJ n = 39). **C**. Table showing the Fischer’s Exact Test comparisons between 5- and 9-month normal circadian and CJ mice. **D**. Representative 200x H&E image from a CJ mouse demonstrating PanIN-1 (arrow) and PanIN-2 lesion (arrowhead). **E**. Representative 200x H&E image from a CJ mouse demonstrating a PanIN-3 lesion (arrowhead) and PDAC off the same gland (dashed circle). Scale bars are shown for reference.

As expected, the incidence of advanced lesions increased by 9 months of age in KC mice under normal circadian conditions to 3% (n = 1/33) harboring PanIN-2, 6.1% (n = 2/33) developing PanIN-3, and 15.2% (n = 5/33) of mice progressing to PDAC (**Figure 2B/C**). Notably, although the incidence of PanIN-3 (10.3% [n = 4/39]) and PDAC (10.3% [n = 4/39]) was similar between CJ and normal circadian KC mice at 9 months, there was a significant increase in PanIN-2 formation in KC mice due to CJ (23.1% [n = 9/39] vs 3% [n = 1/33], *p* = 0.02) (**Figure 2B-E**). There were no sex-specific differences in PanIN-2 incidence for normal circadian (male 0/19 (0%) vs. female 1/14 (7.1%); *p* = 0.42) and CJ mice (male 4/23 (17.4%) vs. female 5/16 (31.2%); *p* = 0.44). Collectively, this data demonstrates that circadian desynchrony promoted by CJ in KC mice caused accelerated development of FI/PanIN at 5 months leading to an increased incidence of higher grade PanINs (i.e. PanIN-2) by 9 months, and these findings were independent of sex. This strongly supports the assertion that circadian misalignment (desynchrony) facilitates tumor initiation and progression of the neoplastic PanIN lesions.

### Pancreatic Clock Gene Deletion Abrogated Tumor Initiation

Different cell populations within the pancreas can lead to tumor development. For instance, islet cells can transform into neuroendocrine tumors (Zhang et al. 2013; Li and Xie 2022), whereas acinar-to-ductal metaplasia is the initiating process in PanIN formation (and consequently PDAC) (Chuvin et al. 2017). Therefore, we hypothesized that misalignment of the clock within acinar cells was responsible for the accelerated PanIN phenotype seen in CJ KC mice. In turn, we employed a cell-autonomous *Bmal1* knock-out model to test whether circadian disruption in acinar cells facilitates tumor initiation and progression of the pancreas cancer precursor lesions. In support of this hypothesis, previous studies in hepatocellular carcinoma (HCC) have revealed a concordant phenotype between CJ and cell-autonomous clock disruption (*Bmal1* knock-out) in mice predisposed to develop HCC (Kettner et al. 2016). We crossed the KC mouse with a mouse harboring a conditional deletion of the core clock gene *Bmal1* to produce *Bmal1*^*fx/fx*^; *LSL-Kras*^G12D/+^; *Pdx1*-*Cre* (BKC) mice, which possess *Bmal1* deletion in cells concomitantly expressing *Pdx1* and the *Kras*^G12D^ mutation (Johnson et al. 2014). BMAL1 is a central transcriptional regulator of the clock and is therefore essential for clock functionality (Bunger et al. 2000). We evaluated the degree of FI/PanIN in BKC mice under standard lighting conditions at 5 months and, contrary to our hypothesis, we found that cell-autonomous clock disruption caused resistance to FI/PanIN development (**Figure 3A-D**). Compared to normal circadian KC mice, BKC mice had scant development of FI/PanIN (12.73% ± 12.08 (KC) vs 1.59% ± 0.65 (BKC); *p* < 0.01), with essentially histologically normal pancreas, and no mice exhibited a higher-grade lesion than PanIN-1 (**Figure 3A-D**). Additionally, this suppressed PanIN development in BKC mice was not sex-dependent (male 1.64% ± 0.74 vs. female 1.52% ± 0.53; *p* =0.65).

**Figure 3:**
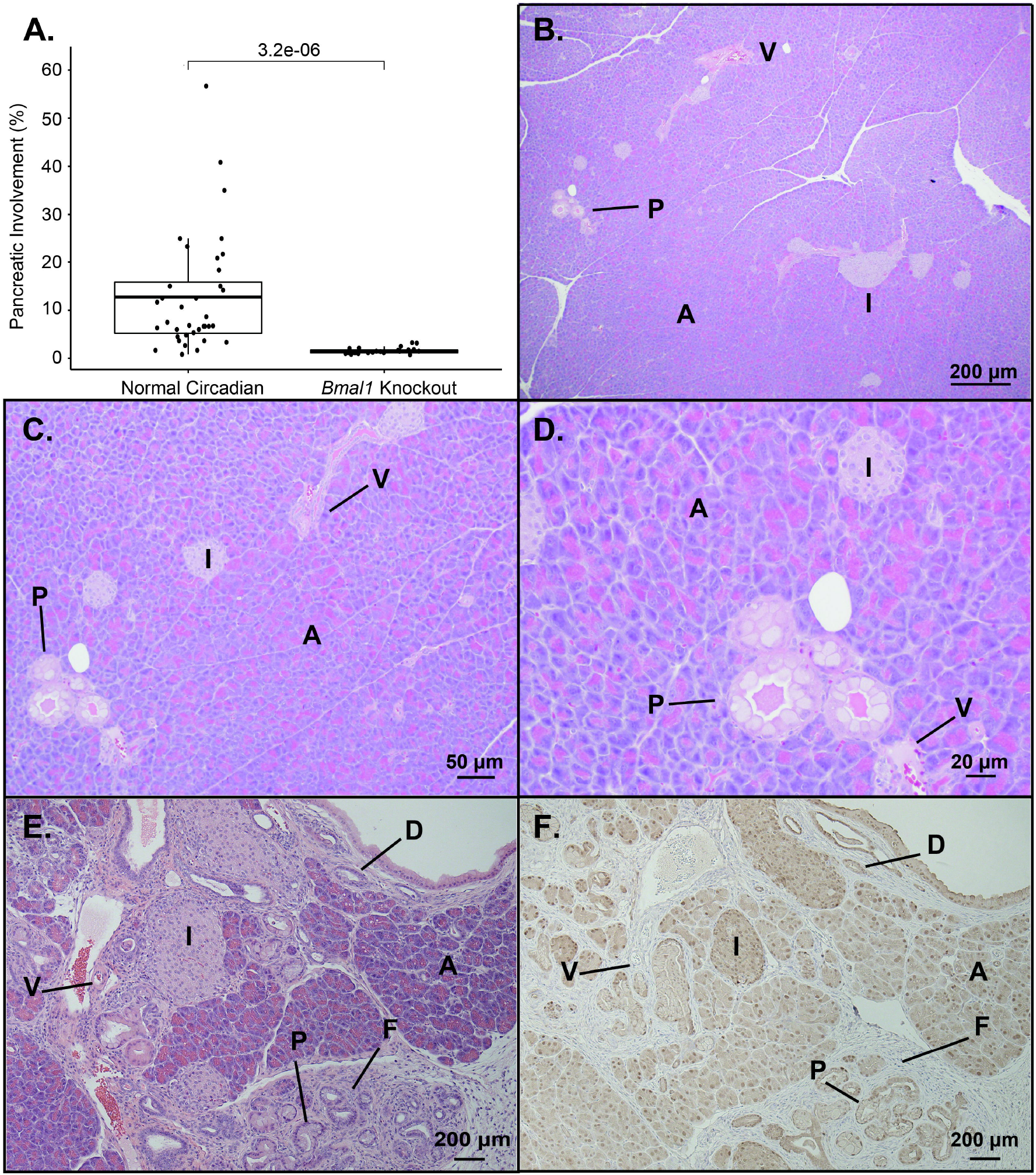
Pancreatic cell-autonomous deletion of Bmal1 does not accelerate pancreatic fibroinflammatory infiltrate/PanIN development at 5 months. **A**. Boxplot comparing the mean (with associated 25^th^ and 75^th^ quantiles) percent pancreatic involvement of FI/PanIN at 5 months for normal circadian KC (n= 36) and BKC mice (n = 24). Representative H&E images demonstrating the FI/PanIN from a BKC mouse at 40x (**B**), 100x (**C**), and 200x (**D**) To evaluate the cell types affected in BKC, KC mice were crossed with Ai14 reporter mice to create KCT mice. H&E stained sections (**E**) and immunohistochemistry for tdTomato (**F**) were compared to evaluate cell-specific Pdx1-driven Cre expression in KC mice. Scale bars are shown for reference. [annotation: PanIN (P), acinar (A) cells, islet (I) cells, ductal (D) cells, vascular (V) cells (i.e. endothelial cells), and fibroinflammatory (F) infiltrate]

The observed results with cell-autonomous clock disruption were discordant with our expectations and demonstrated that *Bmal1* knock-out in the targeted cells did not recapitulate the accelerated PanIN phenotype seen with KC mice under (global) CJ conditions. *Pdx1* is expressed early during pancreatic development and later to commit pancreatic progenitor cells to islets (Offield et al. 1996; Hingorani et al. 2003). To understand which cell populations in BKC mice exhibited loss of BMAL1 (location of *Pdx1* expression), we crossed the KC mouse with the Ai14 ‘marker’ mouse that conditionally expresses tdTomato in the presence of *Pdx1*-driven Cre-recombinase (**Figure 3E-F**). This revealed tdTomato expression (*Kras* mutation and *Bmal1* deletion) in acinar cells, ductal cells, PanINs, and islet cells. Concomitantly, expression was absent in the fibroinflammatory infiltrate (fibroblasts, collagen, and immune cells) and blood vessels surrounding the PanINs. Thus, in mice predisposed to pancreas neoplasm formation, circadian disruption in acinar cells produced a different phenotype compared to CJ, suggesting a different cell population may be responsible for the observed accelerated PanIN formation and progression in CJ KC mice.

### Single-cell RNA Sequencing of KC Mice Recapitulates the Highly Heterogeneous Tumor Microenvironment

Prior work in the liver revealed that cell-autonomous knock-out of the clock in liver hepatocytes (Albumin*-*Cre; *Bmal1*^fx/fx^) drove a nearly identical phenotype compared to CJ mice (non-alcoholic fatty liver disease progressing to HCC) (Kettner et al. 2016). However, we found that in the pancreas of KC mice this was not evident. We expected that this was a consequence of non-cell autonomous changes to the circadian clock (i.e. not acinar cells or PanINs) – whether by loss of clock function or by misalignment – are responsible for accelerating the neoplastic progression. To examine further, we tested the hypothesis that misalignment of the clock in an alternative cell population caused the accelerated PanIN formation and progression; this was done by evaluating the expression of individual cells in the tumor microenvironment over time. We performed single-cell RNA sequencing (scRNAseq) from pooled pancreatic samples (1 male/female) from both normal circadian and CJ KC mice at 5 and 9 months (**Figure 4A/B**). Notably, male and female mice were pooled because we consistently demonstrated no sex-specific effect within either the normal circadian or the CJ group of KC mice at 5 months or 9 months. After filtering and dimensionality reduction, a total of 14 different heterogeneous cell types were annotated using the ScType package comprising 42,211 cells (**Figure 4A-B & Supplemental Data 1-2**) (Ianevski et al. 2022). These included acinar cells, several classes of immune cells, endothelial cells, and fibroblast populations.

**Figure 4:**
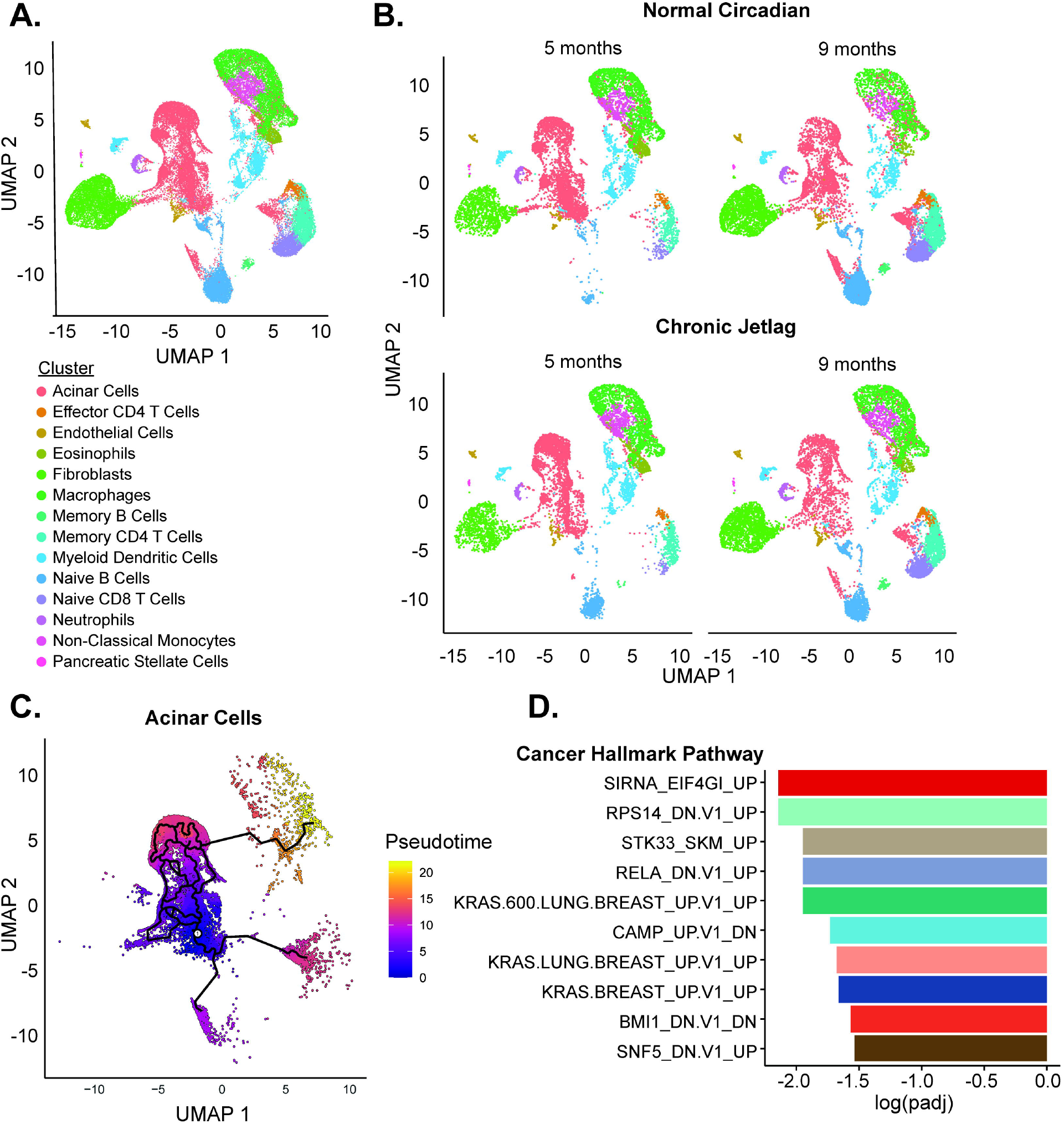
The single-cell landscape of KC mice recapitulates the developing pancreatic cancer tumor microenvironment. **A**. UMAP demonstrating the 14 diverse cell types and 42,211 cells in the scRNAseq dataset. **B**. UMAPs split by time (5 and 9 months) and condition (normal circadian and CJ) **C**. Pseudotime trajectory analysis was performed for the subset of pancreatic acinar cells with 0 pseudotime set to the point with the highest fraction of cells at 5 months. **D**. Gene Set Enrichment Analysis (GSEA) was performed for the oncogenic signature gene set associated with pseudotime progression and ordered by the log adjusted p value.

We first began by subsetting the pancreatic epithelial cells, which should encompass both normal acinar and developing PanINs, and used pseudotime trajectory analysis to track cell-state changes that occurred from 5 to 9 months. The expectation was that this population would gain signatures of neoplastic transformation over time, given the loss of normal acinar cell density and increase in PanINs from 5 to 9 months. Pseudotime was set to zero at the point with the highest fraction of cells from the 5-month mice and allowed to progress (**Figure 4C**). Moran’s I autocorrelation analysis was then performed to obtain genes that varied across the trajectory (**Supplemental Data 3**). Gene set enrichment analysis (GSEA) of the genes that correlated with pseudotime progression revealed an association with multiple *Kras*-driven cancer cell line signatures – corresponding precisely to the development of *Kras*-driven pancreatic neoplasia over time (**Figure 4D**). Therefore, we identified a collection of pancreatic epithelial cells that appeared to represent the transition from acinar cells to PanIN, coinciding with the increased PanIN formation seen in histopathology from 5 to 9 months. These findings established a foundation for the subsequent analysis of the clock to understand the effects of circadian misalignment (CJ) in the acinar cell and PanIN population, which would then help to support or refute our new hypothesis of a non-cell-autonomous cell population driving the accelerated PanIN phenotype.

### Chronic Jetlag Promotes Differential Clock Gene Expression in Fibroblasts

We performed differential gene expression comparing the collective 5- and 9-month normal circadian and CJ mouse cells, and assessed for differences in the CCG and clock-associated genes (including *Tef, Hlf, Bhlhe40, Bhlhe41, Nfil3, Rorc, Dbp*, and *Ciart*) (**Supplemental Data 4**). We previously showed that CJ-induced clock misalignment in the pancreas caused differential expression of the core clock genes (Schwartz et al. 2021), and thus we expected populations of cells manifesting misalignment of the clock between normal circadian and CJ conditions would similarly demonstrate CCG differential expression. In support of our new hypothesis, the acinar cell/PanIN clusters that we identified and confirmed from the pseudotime analysis did not exhibit differential expression of any of the CCGs or the clock-associated genes between clusters derived from normal circadian or CJ KC mice. We then evaluated the other clusters, and we found that endothelial cells, macrophages, memory CD4 T cells, and monocytes each had a single clock-associated gene differentially expressed (**Figure 5A**). Interestingly, fibroblasts were the only cell type that exhibited differential expression of CCGs and clock-associated genes as a result of circadian misalignment, including *Per2, Per3, Nr1d2, Hlf, Tef*, and *Dbp* (**Figure 5A-B**). Fibroblasts are an integral part of the PDAC microenvironment and are known to exhibit circadian rhythmicity when compared to naïve fibroblasts (Parascandolo et al. 2020). The lack of other cell types exhibiting changes in clock-specific gene expression suggests fibroblasts may be the target of the CJ-induced circadian misalignment and accelerated neoplastic phenotype. Concordantly, fibroblasts were among the population of cells that were negative for Pdx1 expression (in the KCT ‘marker’ mouse), corresponding to an intact clock function in the BKC mice.

**Figure 5:**
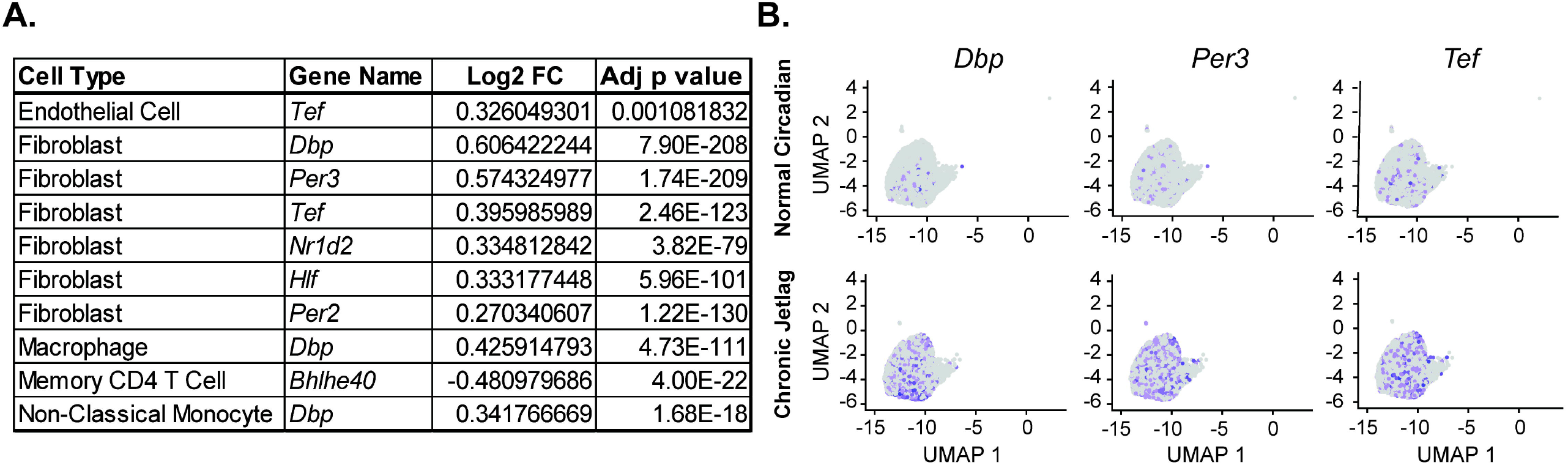
Fibroblasts were preferentially affected by chronic jetlag. **A**. Table of core clock genes (CCGs) and clock-associated genes differentially expressed when comparing normal circadian and CJ conditions of each cell type. **B**. Representative UMAP dimensionality plots demonstrating cellular expression (blue color) of select CCG and clock-associated genes in the subset of fibroblasts.

### Fibroblasts Exhibit Time-Dependent Effects Secondary to Chronic Jetlag

To further evaluate the fibroblasts within our dataset, we isolated three different subclusters (**Figure 6A**). Using existing markers generated for cancer-associated fibroblasts (CAFs), including inflammatory CAFs (iCAFs), myofibroblast CAFs (mCAFs), and antigen-presenting CAFs (apCAFs), we assigned identities with ScType (**Supplementary Data 5-6**) (Elyada et al. 2019). Of the three clusters, we identified all three as iCAFs – with the expression of known markers, such as *Cle3b, Il6, Has1*, and *Col14a1* (**Figure 6B**).

**Figure 6:**
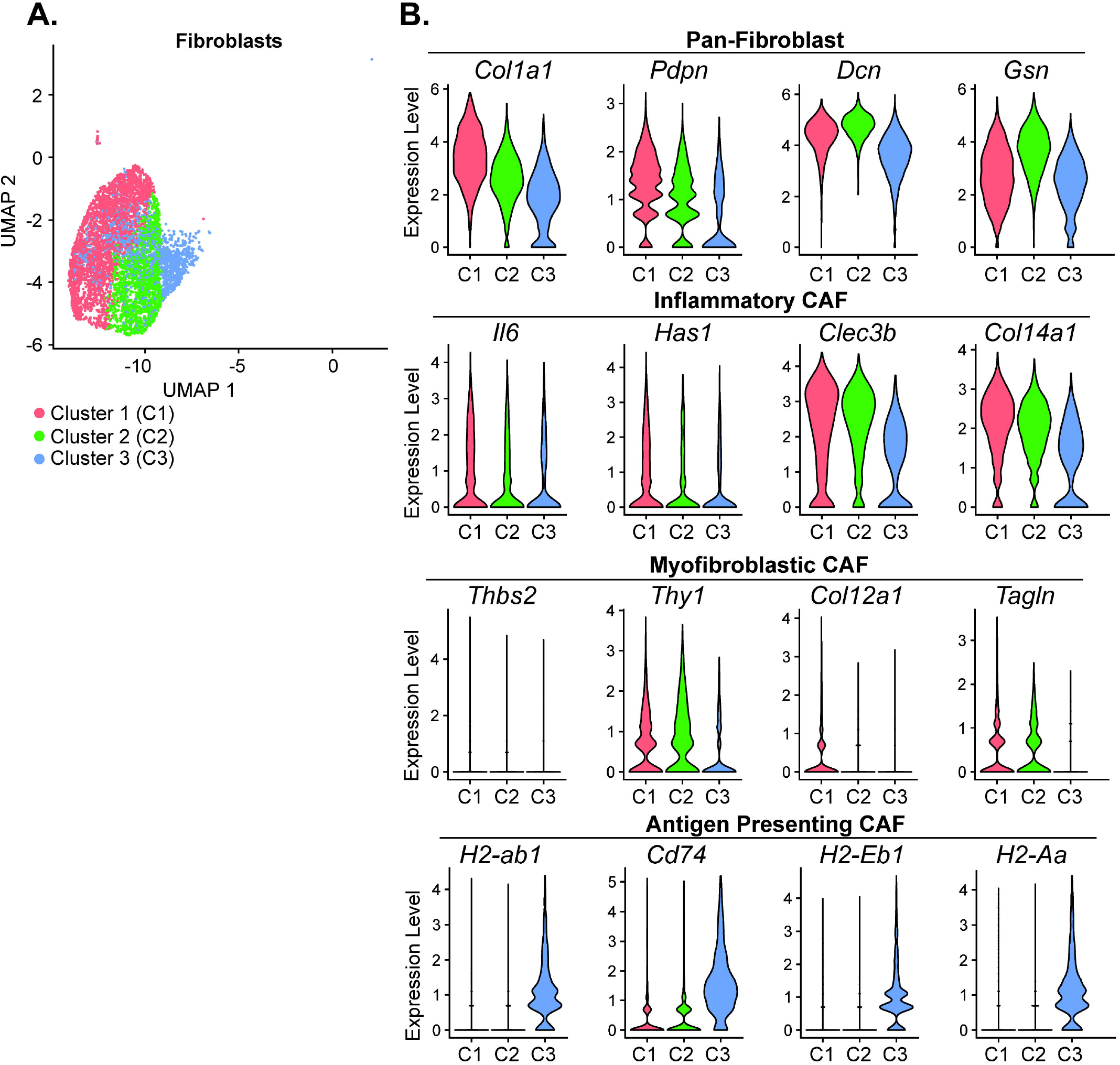
Fibroblasts subclusters in the developing pancreatic tumor microenvironment were identified as inflammatory cancer-associated fibroblasts. **A**. A UMAP demonstrating the three fibroblast subclusters identified on dimensionality reduction. **B**. Violin plots with select markers for Pan-, Inflammatory-, Myofibroblastic-, and Antigen Presenting-Cancer Associated Fibroblasts (CAFs).

Fibroblast cluster 3 exhibited some characteristics consistent with apCAFs, including expression of multiple major histocompatibility complex (MHC) II components *H2-Aa, H2-Eb1*, and *H2-ab1* and *Cd74*, the invariant chain of MHC II (Figure 6B). We then performed differential gene expression and GSEA comparing the iCAFs between normal circadian and CJ groups at 5 and 9 months (**Figure 7A-B** & **Supplemental Data 7**). We found that multiple metabolic pathways in fibroblasts were enriched in the CJ group at 5 months (**Figure 7A**), and our 9-month comparison revealed enrichment of several apoptosis-related pathways, including ‘NEGATIVE_REGULATION_OF_CELL_DEATH’ and ‘REGULATION_OF_APOPTOTIC_SIGNALING_PATHWAY’ (**Figure 7B**). To evaluate if CJ was simply increasing fibrosis (i.e. collagen deposition) to facilitate these changes, we compared the collagen content from 5- and 9-month normal circadian and CJ KC mice (**Supplemental Data 8**). We found no differences in the collagen content at either time point. We, therefore, identified that the iCAFs in the developing tumor microenvironment of KC mice had early enrichment of metabolic pathways in response to CJ, with late enrichment of apoptosis-related pathways. Enrichment of these pathways in fibroblasts as a consequence of CJ corresponds with the finding that fibroblasts manifest differential expression of core clock genes (i.e. affected by misalignment of the clock). When combined with the observation that acinar cells/PanINs demonstrated no differential gene expression of CCGs, and abolishing the clock in acinar cells did not replicate the CJ-induced phenotype, these results suggest CJ causes clock desynchrony in the fibroblast population to alter signaling and accelerate the formation and progression of PanINs.

**Figure 7:**
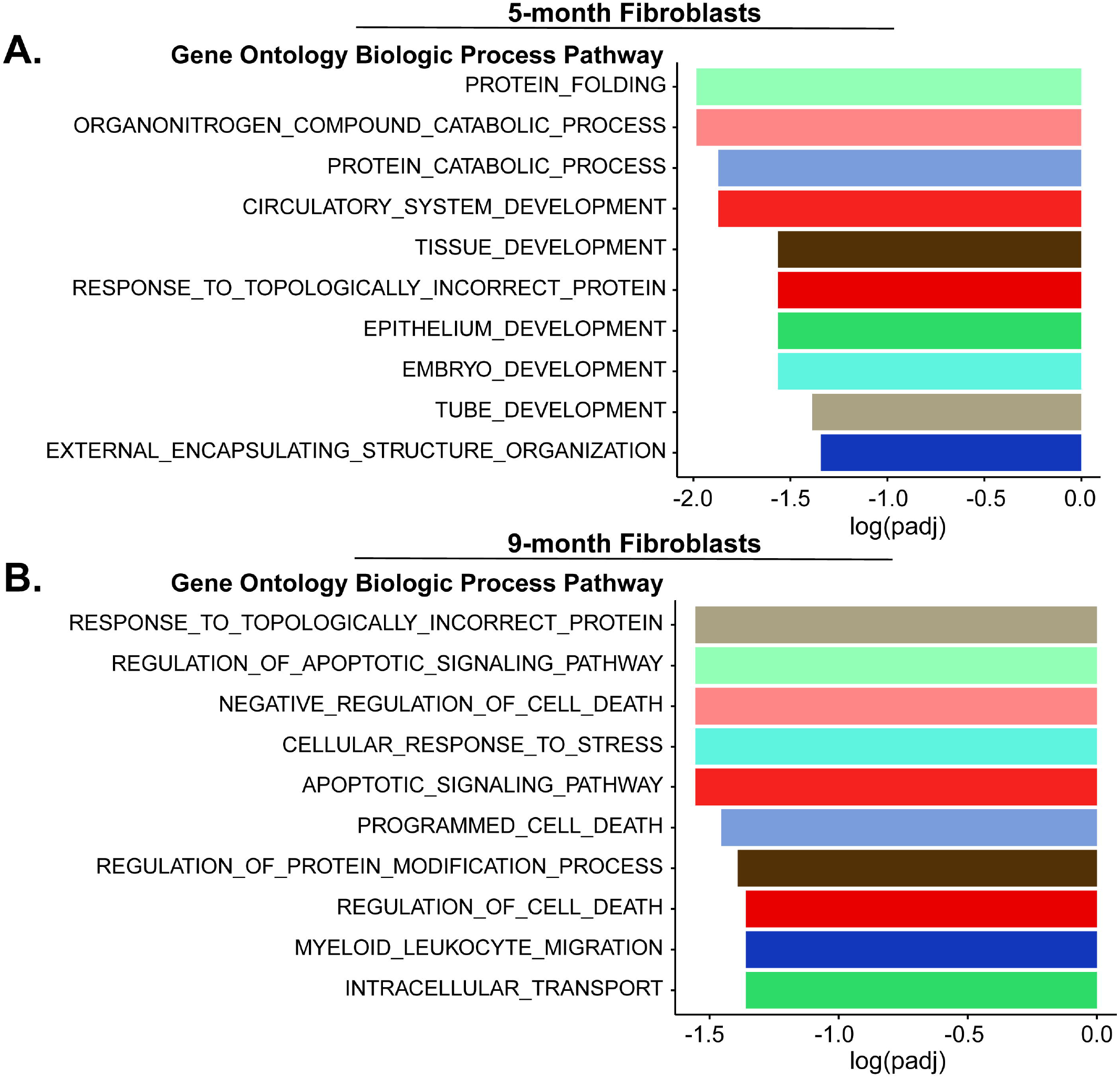
Chronic jetlag influences fibroblast expression patterns across time. Figures with The Gene Set Enrichment Analysis (GSEA) demonstrating the gene ontology biologic process (GO BP) pathways enriched in CJ compared to normal circadian fibroblasts at **A**. five and **B**. nine months. The top 10 pathways at each time point are ordered by their log-adjusted p value.

## Discussion

There has been a substantial volume of literature examining circadian desynchrony, which encompasses the misalignment between environmental or behavioral cycles and the endogenous clock (Vetter 2020). Persistent misalignment between environmental cues (light/dark cycle) and behavioral cues (feeding) compared to the internal clocks can result in aberrant gene signaling (Kettner et al. 2016; Sulli et al. 2018) with consequent disease pathology (Scheer et al. 2009; Lee et al. 2013; Morris et al. 2015). For instance, in healthy human adults subjected to a forced circadian desynchrony protocol that replicated shift work (12-hour reversal of light/dark cycle for 3 days), there was a misalignment of the circadian clock relative to the timing of the light/dark cycle causing glucose intolerance as well as reduced insulin sensitivity – findings indicative of impaired glucose metabolism that occurs with the onset of type II diabetes (Morris et al. 2015). This coincides with epidemiological findings of an increased risk of type II diabetes with shift work (Pan et al. 2011) and is congruent with the increased risk of obesity and metabolic syndrome in individuals performing rotating shift work schedules (Lin et al. 2009; Antunes et al. 2010). Unfortunately, the metabolic reprogramming and chronic inflammation that ensues following these diagnoses can drive the formation of several cancers such as hepatocellular carcinoma (Campbell et al. 2016), which commences a path of ‘neoplastic change’ consisting of non-alcoholic fatty liver disease, non-alcoholic steatohepatitis, and liver cancer (Simon et al. 2021). The same process of neoplastic transformation in the liver can occur due to circadian misalignment: C57Bl/6J mice subjected to chronic jetlag conditions experienced an acceleration of non-alcoholic fatty liver disease and an increased incidence of hepatocellular carcinoma compared to mice under standard lighting conditions (Kettner et al. 2016).

Notably, the evidence correlating circadian desynchrony and liver cancer is mirrored in a similar metabolic organ – the pancreas. There is a strong correlation between obesity and diabetes, and risk of developing pancreatic ductal adenocarcinoma (PDAC). A potential link between circadian misalignment (associated with obesity and diabetes) and PDAC can be inferred from this correlation, and this putative link is strengthened by epidemiologic data reporting an increased risk of PDAC in shift workers (Parent et al. 2012). Therefore, based on this cumulative evidence, we sought to test the hypothesis that circadian clock desynchrony through chronic jetlag (CJ) would accelerate the initiation and progression of PanINs in mice predisposed to develop pre-cancerous lesions (PanIN-1,-2,-3) and PDAC.

Our work revealed that circadian misalignment through CJ increased tumor initiation and progression of PanINs, which are pancreas cancer pre-cursor lesions that can ultimately develop into PDAC. Our findings of accelerated neoplasm progression in the current study are alarming considering the highly lethal nature of PDAC (Siegel et al. 2021), and the prevalence of circadian desynchrony in society (Antunes et al. 2010). This type of clock dysfunction affects a significant percentage of the population, such as shift workers under imposed non-daytime schedules, who experience chronic misalignment between their endogenous clock and environmental cues (Bass and Lazar 2016). A less severe but highly prevalent form of misalignment may also be present through extended exposure to light at night, extended duration of dietary intake, or significant alteration in sleep patterns (social jetlag) (Roenneberg et al. 2012; Chang et al. 2015; Bass and Lazar 2016; Fishbein et al. 2021). To study the potential implications of these types of clock desynchrony in the development of PDAC, we utilized a mouse model in which *Kras*-mutation in pancreatic acinar cells promotes acinar-to-ductal metaplasia, followed by PanIN formation, and ultimately PDAC. Consequently, a pertinent question for the present work is the applicability of the model to humans. Interestingly, in humans, PanINs are frequently found in the pancreas and their prevalence increases with age (Matsuda et al. 2017). They are a result of acinar metaplastic transformation to a ductal cell phenotype, typically in response to injury or inflammation (De La O et al. 2008; Reichert and Rustgi 2011; Chuvin et al. 2017). In an autopsy study of 173 cases with no evidence of PDAC (mean age 80.5 years), PanIN-1 was present in 77% of pancreas specimens, PanIN-2 was present in 28%, and 4% of the pancreas specimens harbored PanIN-3 (Matsuda et al. 2017).

Consistent with our KC model, PanIN-1s were always present in the pancreas when PanIN-2 or -3 were identified. These autopsy findings were also confirmed in a separate study (Longnecker and Suriawinata 2022). Thus, our model is concordant with the neoplastic process that occurs in humans and relevant for studying PanIN progression. Our findings of CJ-induced acceleration of advanced PanIN formation could be applicable to many individuals who unknowingly harbor PanINs and experience consistent circadian misalignment.

PanINs are microscopic and early PanINs are unable to be detected with the current imaging modalities (Kanda et al. 2012). Therefore, human studies of circadian misalignment driving PanIN progression have not been feasible. Yet, the *in vivo* findings in our study warrant consideration of future investigations designed to evaluate risk in humans and possibly risk mitigation. Given the widespread presence of circadian misalignment, and the prevalence of PanINs, a suitable population of individuals to target for risk mitigation of circadian desynchrony may prove tenable. For example, there is convincing evidence that patients who have a family history of PDAC demonstrate a greater propensity for PanIN formation, are more likely to harbor advanced PanINs, and, therefore, are at higher risk for developing PDAC. In a study by Shi and colleagues evaluating patients who underwent pancreatic resection for PDAC, those with a family history of PDAC (without a germline mutation) demonstrated a rate of 1.51 PanIN lesions/cm^2^ in the pancreas (i.e. non-cancer pancreas) compared to 0.55 PanIN lesion/cm^2^ in patients with sporadic PDAC (Shi et al. 2009). The majority of the lesions were PanIN-1, with fewer PanIN-2 and PanIN-3 lesions identified (Shi et al. 2009); again mirroring the findings in our KC model where there is multifocal lesion formation, and the majority of the lesions at 9 months were PanIN-1 with fewer PanIN-2 and PanIN-3. Moreover, in the study by Shi *et al* (Shi et al. 2009), PanIN lesions were found in the pancreas remote from invasive cancer, which supports the findings from autopsy studies (Matsuda et al. 2017; Longnecker and Suriawinata 2022) that not all PanINs progress to cancer. Similar to the etiology of PanINs in our model, greater than 95% of human PanINs harbor a *Kras* mutation (including 92% of PanIN-1) (Kanda et al. 2012), and it is generally accepted that the fraction of *Kras*-mutant cells in PanINs increases with PanIN grade, indicating clonal expansion of the *Kras*-mutant cells that have been ‘transformed’ (Kanda et al. 2012). However, the drivers of this transformation from PanIN-1 to higher-grade PanINs or PDAC are poorly understood. Consequently, our study forms a strong basis for further investigation to understand how circadian misalignment promotes the progression of PanIN lesions. This should lead to prevention strategies (e.g. chemoprevention or behavioral modification) that can then be applied to high-risk patients to prevent the progression of PanINs to cancer (Miller et al. 2016). Incidentally, these high-risk patients are already being captured in high-risk clinics, where individuals with a strong family history or known pre-cursor lesions in the pancreas are being followed to identify cancer transformation at an early stage (Shin et al. 2015; Miller et al. 2016; Dbouk et al. 2022). This would be an ideal group to target for risk mitigation or chemoprevention.

To better understand the basis for our observations of accelerated neoplasm formation with circadian misalignment (CJ), we implemented a conditional mouse model to achieve cell-autonomous core clock gene knock-out. We chose a conditional knock-out as opposed to a global knock-out because of the known difficulty in disentangling the effects in a global knock-out model (e.g. *Bmal1*^*-/-*^). For example, the opposing effects in glucose metabolism and insulin sensitivity with knock-out of the clock in the pancreas and liver can mask glucose intolerance phenotypes that are otherwise readily apparent with islet-cell-specific clock disruption (Lamia et al. 2008; Marcheva et al. 2010; Perelis et al. 2016). Furthermore, chronic jetlag exposure is a ‘global’ effect and we aimed to home in on the cell population responsible for the observed phenotype. In this way, the use of core clock gene knock-out models has been indispensable in ascertaining clock-dependent mechanisms for various pathologies. Oftentimes, the phenotypes among circadian disrupted groups are concordant, whether by circadian desynchrony or through genetic alteration. For instance, Lee and colleagues found that the cell-autonomous deletion of *Bmal1* in β-cells of the pancreas resulted in the loss of glucose-stimulated insulin secretion, and the blunted insulin secretion with peripheral insulin resistance was replicated by environmentally-induced circadian misalignment (i.e. 6-hr light phase advancement protocol) (Lee et al. 2013). In a separate study, mice predisposed to colorectal cancer development were subjected to circadian misalignment via constant lighting conditions and showed a 2-fold increase in tumor initiation compared to those housed under diurnal conditions (Stokes et al. 2021). In this study, Stokes *et al*. showed that global and epithelial loss of *Bmal1* similarly induced tumor initiation, suggesting that *Bmal1* expression in the colon epithelial cells (cell of origin for colorectal cancer) is similarly ‘targeted’ by perturbations in the clock (altered lighting) and by clock disruption (*Bmal1* deletion). Comparable results were seen by Kettner *et al*. when evaluating rates of non-alcoholic fatty liver disease and hepatocellular carcinoma between cell-autonomous *Bmal1* knock-out mice and mice subjected to circadian desynchrony (Kettner et al. 2016). Thus, we expected that cell-autonomous clock disruption in KC mice (pancreatic endocrine and exocrine *Bmal1* knock-out) would demonstrate a PanIN phenotype concordant with the circadian misaligned KC mice. However, when we crossed our KC mice with the condition *Bmal1*^*fx/fx*^ mouse, we found a discordant phenotype in pancreatic tumor initiation compared to circadian misalignment. Although the results were unanticipated, they did provide insight and enabled us to conclude that disruption of the clock in acinar cells does not facilitate PanIN formation or progression. This led us to investigate separate putatively involved cell populations.

To better understand individual cell population involvement, and to build a foundation for discerning how pancreatic neoplasia more rapidly progressed in CJ conditions, we performed scRNAseq at both 5 and 9 months. We found that the pancreatic epithelial cells, including the PanINs and acinar cells, showed a time-dependent cancer signature on pseudotime analysis consistent with the observed histopathologic changes. Confirming the time-dependent cancer signature enabled us to examine these cells primarily for changes to the clock, to determine if the scRNAseq data from circadian misaligned mice complemented the *Bmal1* cell-autonomous knock-out data. Subsequently, we assessed 13 other cell types for evidence of differential gene expression between normal circadian and CJ, and we found that fibroblasts –not the pancreatic epithelial cells – were the only cell type with multiple CCG and clock-associated gene changes. This finding supports the notion that CJ augments fibroblasts in the developing tumor microenvironment and not necessarily the epithelial/PanINs themselves. Notably, this also reinforces the differences identified in the cell-autonomous *Bmal1* knock-out model versus the circadian desynchrony model – misalignment may be targeting the fibroblast population to accelerate neoplasm formation and progression. Meanwhile, in the *LSL-Bmal1*^*fx/fx*^; *Pdx1-Cre; LSL-Kras*^*G12D/+*^ mice, *Bmal1* (and thus the clock) is intact in the mesenchymal cells: IHC in KC mice conditionally expressing tdTomato demonstrated that there was *Pdx1-*Cre-directed expression in the acinar, ductal, islet, and PanINs, but not the mesenchymal lineage cells, including the fibroblasts and endothelial cells. We used additional expression markers to subset the fibroblasts and we identified all 3 clusters as inflammatory CAFs (iCAFs) based on established markers (Elyada et al. 2019). iCAFs secrete IL-6, CXCL9, and TGFβ and can be both immunosuppressive as well as immune-promoting (Sahai et al. 2020). Interestingly, we did not observe any differences in the amount of pancreatic collagen content at 5 or 9 months between normal circadian and CJ KC mice, suggesting a role for paracrine signaling in mediating the effects of CJ by fibroblasts rather than extracellular matrix production (i.e. desmoplasia). While this requires additional consequential investigation and will be the focus of future work, it is important to note that other investigators have uncovered the substantive involvement of the fibroblast clock in cancer. In a pan-cancer analysis of The Cancer Genome Atlas tumor samples, Wu and colleagues employed a new computational approach (LTM) to explore clock-controlled pathways in cancer (Wu et al. 2022). The authors found that the variation in clock strength across 9 separate cancers was directly related to the proportion of fibroblasts in the samples and concluded that this was consequent to fibroblasts possessing a more robust circadian clock than the adjacent cancer cells within the tumor microenvironment. Moreover, fibroblasts are known to have an essential role in fostering cancer progression in the pancreas cancer microenvironment (Hanahan and Coussens 2012; Helms et al. 2020). As fibroblast density increases with the advancement of PanINs (i.e. PanIN-1 to PanIN-2/3) towards invasive cancer, the cancer-associated fibroblasts (CAFs) display a ‘wound-healing’ type response that supports neoplastic proliferation through direct and indirect metabolic contributions (Helms et al. 2020). CAFs display differences in rhythmic gene expression compared to naïve fibroblasts (Parascandolo et al. 2020), and the wound healing type response in fibroblasts has been previously shown to be strongly clock dependent (Hoyle et al. 2017), with disruption of the clock profoundly affecting this pathway. Concordantly, after identifying and characterizing the CAFs in our mouse pancreas samples, we performed GSEA at 5 and 9 months. This demonstrated that at 5 months CJ promoted metabolic gene enrichment in the CAFs, and by 9 months, there was enrichment of multiple cell death and apoptosis-related pathways. It is therefore conceivable that the fibroblasts – which harbor a strong clock and coevolve with PanINs – underlie the circadian desynchrony-induced acceleration of PanIN formation and progression. To investigate further whether circadian disruption in the fibroblasts may recapitulate the circadian misalignment (CJ) phenotype, in future work we aim to generate *Col1a1-Cre; Bmal1*^*fx/fx*^ in FLP recombinase *Kras*-mutant mice (*Pdx1-FlpO*; *FSF-Kras*^*G12D/+*^) (Wu et al. 2017). This would be effective in abolishing the fibroblast clock in the developing tumor microenvironment because all the identified fibroblast populations expressed high levels of *Col1a1* (scRNAseq).

A limitation to consider from the scRNAseq data was that we did not identify islet cell clusters and were therefore unable to discern whether differential expression of CCGs was present in this population of cells. However, Pdx1 is expressed in endocrine cells in the pancreas, and therefore the BKC mice (which did not display an accelerated PanIN phenotype) would have a disrupted clock in this cell population (Offield et al. 1996; Hingorani et al. 2003). While clock disruption via *Bmal1* knock-out in islet cells produces metabolic changes that could ostensibly alter the formation or growth trajectory of tumors (PanINs), the prevailing phenotype with BMAL1 deficiency in islet cells – and also with circadian misalignment – is glucose intolerance, hypoinsulinemia, and hyperglycemia (i.e. characteristics of diabetes) (Marcheva et al. 2010; Lee et al. 2013; Perelis et al. 2016). Given that diabetes is a risk factor for PDAC development (Abbruzzese et al. 2018), and the lack of phenotype (expedited PanIN formation) in the BKC mice, it seems unlikely that circadian misalignment in the islets would have contributed to accelerated PanIN formation and progression. Regardless, we were not able to evaluate this cell population in the scRNAseq analysis, and so this remains a limitation. To investigate further, one could consider beta-cell specific *Bmal1* knock-out (*Rip-Cre; Bmal1*^*fx/fx*^) in FLP recombinase *Kras*-mutant mice (*Pdx1-FlpO*; *FSF-Kras*^*G12D/+*^) to determine if the accelerated PanIN development emerges (Wu et al. 2017). Another consideration is the intriguing observation that loss of pancreatic *Bmal1* (acinar cells, ductal cells, islets, PanINs) resulted in near abolishment of PanIN formation, and not just an absence of accelerated PanIN formation. This finding speaks to the complex interplay of the circadian clock amongst cell populations in the tumor microenvironment. In future work, we aim to employ various genetically engineered mouse models to disentangle these findings, which should have clinical relevance for the multitude of individuals that harbor PanINs while concomitantly experiencing chronic circadian misalignment.

Although previous evidence suggests a role for the circadian clock in PDAC growth and spread, this is the first study to show perturbations of the clock increases tumor initiation in PDAC (Jiang et al. 2016; Jiang et al. 2018). Our study provides a foundation for future work to expand our understanding of how a dyssynchronous clock orchestrates cancer development in the pancreas.

## Supplementary Data

<u>Supplemental Data 1</u>: Marker database for single-cell RNA sequencing cell-type identification

<u>Supplemental Data 2</u>: ScType Scores for overall single-cell RNA sequencing dataset

<u>Supplemental Data 3</u>: Moran’s I pseudotime analysis for the pancreatic acinar cells

<u>Supplemental Data 4</u>: Differential gene expression between normal circadian and chronic jetlag conditions for all cell types

<u>Supplemental Data 5</u>: Marker database for fibroblast subtype identification

<u>Supplemental Data 6</u>: ScType Scores for fibroblast subtype identification

<u>Supplemental Data 7</u>: Differential gene expression between normal circadian and chronic jetlag conditions for fibroblasts at 5 and 9 months

<u>Supplemental Data 8</u>: Quantification of collagen content in 5-month normal circadian and chronic jetlag KC mice

## Acknowledgments

We would like to take the opportunity to acknowledge the University of Wisconsin Carbone Cancer Center Support Grant P30 CA014520, which provides funding for the Experimental Animal Pathology Lab Core.

We would also to acknowledge the Gene Expression Center and the Bioinformatics Resource Center (BRC) at the University of Wisconsin, Madison for their contributions to this work.

Finally, we would like to thank the Michael W. Oglesby Foundation for their funding support of our work in circadian disruption and pancreas pathology.

## Data Statement

Single-cell RNA sequencing data is made publicly available through gene expression omnibus (GEO) (Accession number: GSE209629). All other data that support the findings of this study are available from the corresponding author, SRK, upon reasonable request.

