## Supplemental Data 8 for "Chronic Jetlag Accelerates Pancreatic Neoplasia in Conditional *Kras*-Mutant Mice"

**Methodology:** Select unstained slides that were representative of the percent average pancreatic fibroinflammation/PanIN (as determined on adjacent H&E staining) from 5- and 9-month normal circadian and chronic jetlag (n = 6 for each condition) mice were selected for Masson’s Trichrome staining. Trichrome staining was performed by the Translational Research Initiatives in Pathology (TRIP) lab at the University of Wisconsin. Slides were then scanned for analysis with the Aperio Digital Pathology Slide Scanner (Leica Biosystems, Wetzlar, Germany). ImageJ version 1.53c was then used to analyze the percent pancreatic collagen content by dividing collagen content by the total tissue area (Schneider et al. 2012). The percent pancreatic collagen content was then compared between the two conditions with a t-test. All statistical tests were performed with R version 4.2.0 (Vienna, Austria).

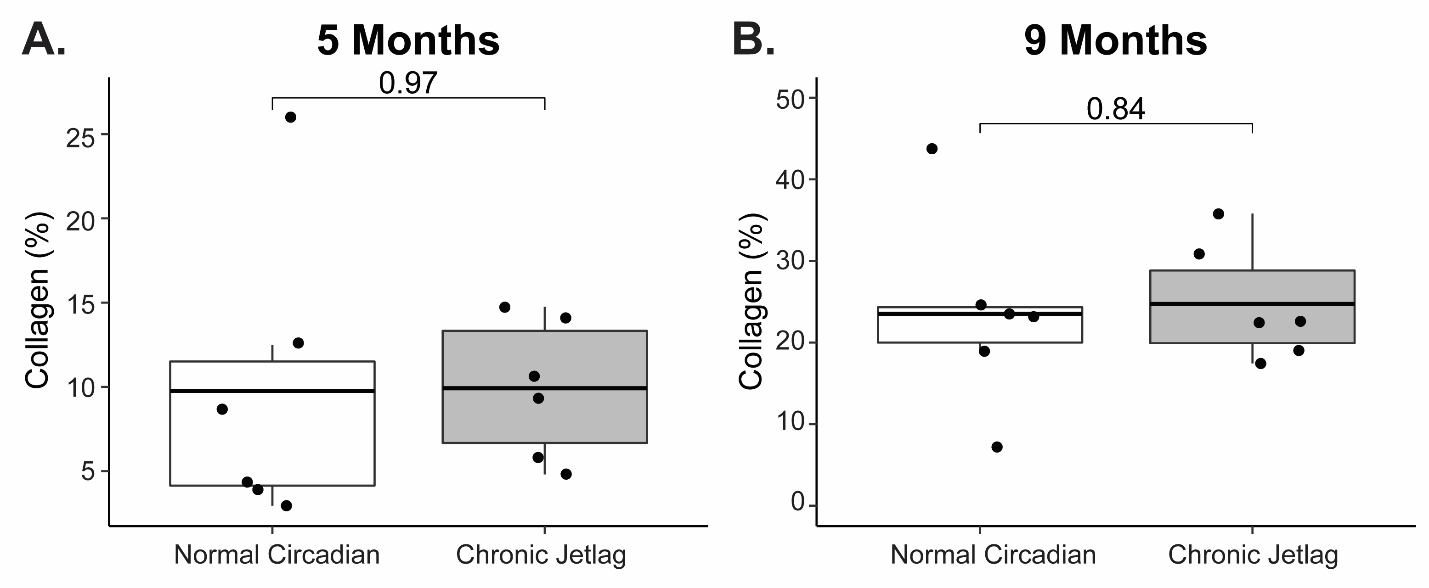

**Figure 1:** Pancreatic c*ollagen content is similar between normal circadian and chronic jetlag KC mice at five and nine months.* **A.** Box plot demonstrating the mean (with associated 25^th^ and 75^th^ quantiles) collagen content at 5 months between normal circadian (n = 6; mean ± se = 9.77% ± 3.57) and CJ (n = 6; 9.92% ± 1.69) KC mouse pancreas (*p* = 0.97). **B.** Similarly, boxplot demonstrating no mean (with associated 25^th^ and 75^th^ quantiles) differences between normal circadian (n = 6; 23.52% ± 4.83) and CJ (n = 6; 24.72% ± 2.92) KC mouse pancreatic collagen content (*p* = 0.84) at 9 months.
